# Cytomove: a browser-local and reviewable workflow for scratch wound healing assay quantification

**DOI:** 10.64898/2026.06.06.730617

**Authors:** Zekeriya Duzgun

## Abstract

The in vitro scratch wound healing assay is one of the most widely used methods for studying collective cell migration, but converting assay images into reproducible measurements remains a practical bottleneck of manual tracing, local software installation, parameter bookkeeping, and limited visibility into how the wound region was segmented. We present Cytomove, a browser-local software tool for reviewable scratch wound healing assay quantification. Cytomove imports local microscopy images, segments the wound region with an explainable variance-and-threshold pipeline implemented in client-side JavaScript without external image-processing dependencies, displays the segmentation as an inspectable overlay before any number is exported, supports single-image and grouped time-course analysis, and exports wound area, wound area fraction, wound width profile statistics, quality-control labels, and full analysis metadata as CSV, Excel, PNG, and ZIP. All processing runs in the browser or in a desktop package built on the same code; microscopy images never leave the user’s machine. In a preliminary comparison with the ImageJ/Fiji Wound Healing Size Tool (WHST) across five image sets and 31 paired measurements, Cytomove reproduced wound-area behaviour closely in a clean brightfield comparator sequence (mean absolute percentage error 4.1%, Pearson r = 0.9975) and in a phase-contrast time course approaching closure (median area error 6.6%, r = 0.9984), while surfacing near-closure and real-world acquisition difficulties through overlays and quality-control labels. Informal local testing indicates that typical single-image analysis completes within seconds in a modern browser, with no installation or dependency step. Cytomove lowers installation friction, keeps assay data local, and links every exported number to the segmentation image and parameters that produced it.

## Introduction

The in vitro scratch wound healing assay is a convenient and inexpensive method for studying collective cell migration. A scratch or gap is introduced into a confluent monolayer of adherent cells, the gap is imaged over time, and the rate at which cells move into the cell-free region serves as a proxy for migratory behaviour. The assay remains popular because it is technically accessible, requires no specialised reagents, is compatible with a wide range of adherent cell types, and produces an intuitive readout of coordinated cell movement (Grada et al., 2017; Jonkman et al., 2014; Liang et al., 2007), a process central to morphogenesis, tissue repair, and cancer invasion (Friedl and Gilmour, 2009). It is used routinely in cancer biology, wound healing, vascular biology, and drug-screening contexts.

The simplicity of the wet-lab procedure contrasts sharply with the difficulty of the image analysis. Manual measurement of wound area or width is slow and subjective, and it scales poorly when an experiment contains many fields, time points, replicates, or treatment arms. General-purpose platforms such as ImageJ and Fiji (Schindelin et al., 2012; Schneider et al., 2012), and dedicated bioimage-analysis environments such as CellProfiler (Carpenter et al., 2006; Stirling et al., 2021) and ilastik (Berg et al., 2019), are powerful and widely adopted, but scratch-assay users must still install plugins or macros, choose parameters, inspect the resulting masks, and keep a record of the settings that produced each measurement. In practice, these settings are frequently lost, which undermines reproducibility even when the underlying numbers are sound.

Several dedicated tools have shaped current expectations for scratch-assay software. TScratch introduced automated wound detection using the fast discrete curvelet transform combined with automated thresholding, paired with visual inspection and optional manual correction; this established that segmentation should remain editable and inspectable rather than hidden (Gebäck et al., 2009). The ImageJ/Fiji Wound Healing Size Tool (WHST) provided a local-variance and thresholding plugin that reports wound area, area fraction, average width, and width variability, and it set a measurement vocabulary that has become a de facto standard (Suarez-Arnedo et al., 2020). PyScratch demonstrated a Python graphical interface for batch and time-course analysis using Laplacian edge detection and Gaussian blurring for non-programmer users, reporting approximately six-fold speed improvement over semi-automated analysis (Garcia-Fossa et al., 2020). CSMA emphasised cell detection within the wound region and two-mode quantification (area and width) using contrast-limited adaptive histogram equalisation and morphological operations (Thanh Pham et al., 2025). Other approaches have combined texture-based segmentation with kinematic readouts, including high-throughput brightfield assays (Zordan et al., 2011) and objective continuous-kinematics curve fitting (Topman et al., 2012).

These tools also leave practical, workflow-oriented gaps: they require local software environments, depend on manual parameter tracking, and are less naturally connected to browser-based sharing, review, and export. The remaining opportunity is therefore not a new biological metric but lower installation friction, clearer provenance, easier group review, explicit linkage between the exported number and the segmentation that produced it, and the ability to keep unpublished microscopy on the researcher’s own machine. Cytomove was developed around this opportunity.

The field is also moving toward more spatially explicit and reproducible assay designs. Recent work has addressed the reproducibility of scratch creation itself through low-cost robotic scratching platforms (Lin et al., 2024), and spatially resolved analysis frameworks now combine wound-edge detection with local cell-trajectory or sector-based measurements (Vašinková et al., 2025). These developments reinforce the same practical requirement: scratch-assay software should make the image evidence, segmentation assumptions, and analysis context visible alongside the number it reports.

## Materials and Methods

### Software implementation

Cytomove is implemented as a single-page browser application using HTML5, the Canvas API, and typed-array operations (Uint8Array, Float32Array, Int32Array) in ES2020+ JavaScript. The entire segmentation pipeline runs client-side; no external image-processing library (ImageJ, OpenCV, or equivalent) is invoked, and no image data is transmitted to a server. A desktop package built with Electron wraps the same analysis code for heavier local workflows and more predictable performance on large images. The public project repository is hosted at https://github.com/zduzgun/CytoMove and a live web version is accessible at https://cytomove.com.

The zero-dependency design means that Cytomove runs in any modern Chromium- or Firefox-based browser without installation or administrator privileges. In informal local testing, typical single-image analysis completed within seconds on a modern consumer laptop or desktop. Group-level operations (loading, previewing, and analysing a time-course set) were usable for sets of up to approximately 50 frames tested in the current implementation, although formal runtime benchmarking across browsers and operating systems remains planned work. The desktop package provides more consistent performance for very large images or long time-course sets by avoiding browser memory limits.

### Segmentation algorithm

The Cytomove segmentation pipeline is an explainable, dependency-free sequence of nine stages. Each stage operates on typed arrays over the raw pixel buffer and corresponds directly to a named step in the exported analysis log, so the contribution of each processing decision remains traceable after the fact.

**Stage 1: Grayscale conversion.** The RGB image is converted to luminance using the ITU-R BT.709 / sRGB perceptual weighting:

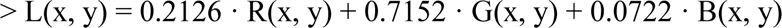

This weighting is perceptually uniform and matches the luminance standard used in display colour spaces; it differs from the older BT.601 coefficients (0.299/0.587/0.114) and from simple channel averaging.

**Stage 2: Field-of-view masking.** An optional cutoff parameter masks dark corners and circular microscope-field borders so that only the imaged field contributes to segmentation. In brightfield mode, pixels below the selected luminance cutoff are excluded from the field mask; for automatic field-of-view cropping, corner luminance samples are also used to estimate dark microscope borders. In phase-contrast (full-field) mode, the field mask covers the full image rectangle. The FOV cutoff is therefore an intensity-threshold control in display-luminance units rather than a percentage.

**Stage 3: Contrast enhancement.** Pixel intensities are clipped to the 1st and 99th percentile values and linearly normalised to [0, 255], stabilising the subsequent variance and threshold steps against illumination outliers and uneven exposure.

**Stage 4: Local variance map.** A local variance is computed for every pixel within a square window of user-selected radius r using a summed-area table (integral image) approach (**Crow, 1984**; **Viola and Jones, 2001**). Given the luminance image I, two summed-area tables are pre-computed:

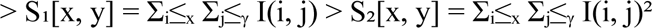

The local mean μ and local variance σ² over the (2r+1)×(2r+1) window centred at (x, y) are then:

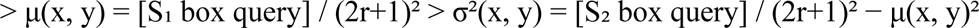

Each box query requires four array lookups, so variance at every pixel is computed in O(1) time regardless of radius. Confluent cell regions exhibit high local texture variance due to cell boundaries, nuclear contrast, and surface texture, whereas the cell-free wound corridor is comparatively smooth and low-variance. This distinction forms the basis for subsequent segmentation.

**Stage 5: Thresholding.** The variance map is thresholded to separate the low-variance wound from the high-variance cell sheet. Otsu’s method (Otsu, 1979) is applied to the variance map to find the threshold T that minimises within-class variance of the binary partition. When the Otsu threshold collapses toward zero on very low-contrast images (a condition detected by comparing T to a percentile-based fallback value), the fallback is applied and the event is recorded in the export log under the field threshold_fallback_used. The user-exposed threshold parameter applies an additive offset to the raw Otsu value, allowing fine adjustment without requiring the user to interpret raw variance units.

**Stage 6: Island filtering.** Connected components in the initial binary map that fall below a user-set minimum size (in pixels squared) are removed before hole filling. This step suppresses isolated debris, speckles, and small non-wound regions that would otherwise persist as noise in the final mask.

**Stage 7: Small-hole filling.** Small enclosed holes inside the wound region are identified using a border flood-fill and inversion procedure, related to the Fill Holes operation in standard binary morphology (Soille, 2004). The selected island-handling level sets the maximum internal hole area that is filled. Larger internal islands are preserved and reported for visual review, so this step suppresses small speckles without hiding substantial cell clusters or bridges inside the wound gap.

**Stage 8: Continuity and component selection.** Connected components are scored by a combination of central proximity, axis span, and area, and the wound-like continuous component is retained. Additional post-processing steps are applied depending on the selected microscope mode: fragmented components within the wound corridor are bridged; a stable wound corridor is extended to the image frame edge where the wound continues beyond the field of view; and, for phase-contrast images, narrow internal slit artefacts are closed using morphological operations. The bridge, edge-extension, and slit-close pixel counts are recorded in the export log.

**Stage 9: Width estimation.** After applying the selected scratch orientation and any fine rotation, the horizontal span of the wound mask is measured at each image row. Cytomove reports the mean, median, standard deviation, and coefficient of variation of the row-wise widths, together with the count and fraction of image rows classified as valid (i.e., rows within the field of view where the wound is detected). The full width profile supports time-course quality-control summaries and frame-level review labels.

### Algorithm parameters and presets

Table 1 lists all user-adjustable segmentation parameters, their ranges, and the values used by each image-type preset. Presets set sensible defaults for common acquisition contexts; any parameter can be overridden individually. When a group of images is loaded, Cytomove samples the images and selects an initial preset automatically; manual preset or microscope-mode changes are treated as user overrides and are not overwritten by subsequent auto-detection.

**Table 1.**
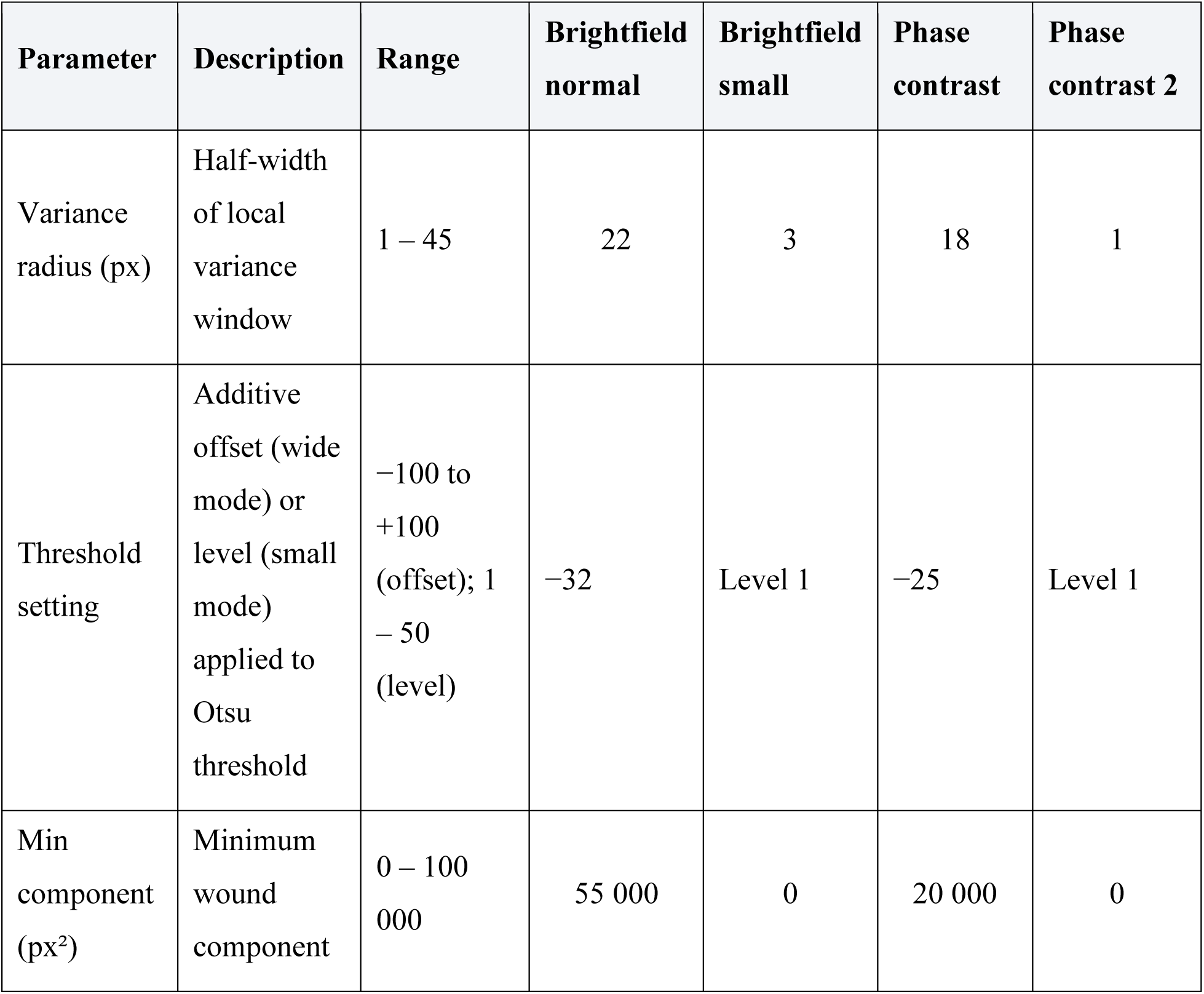
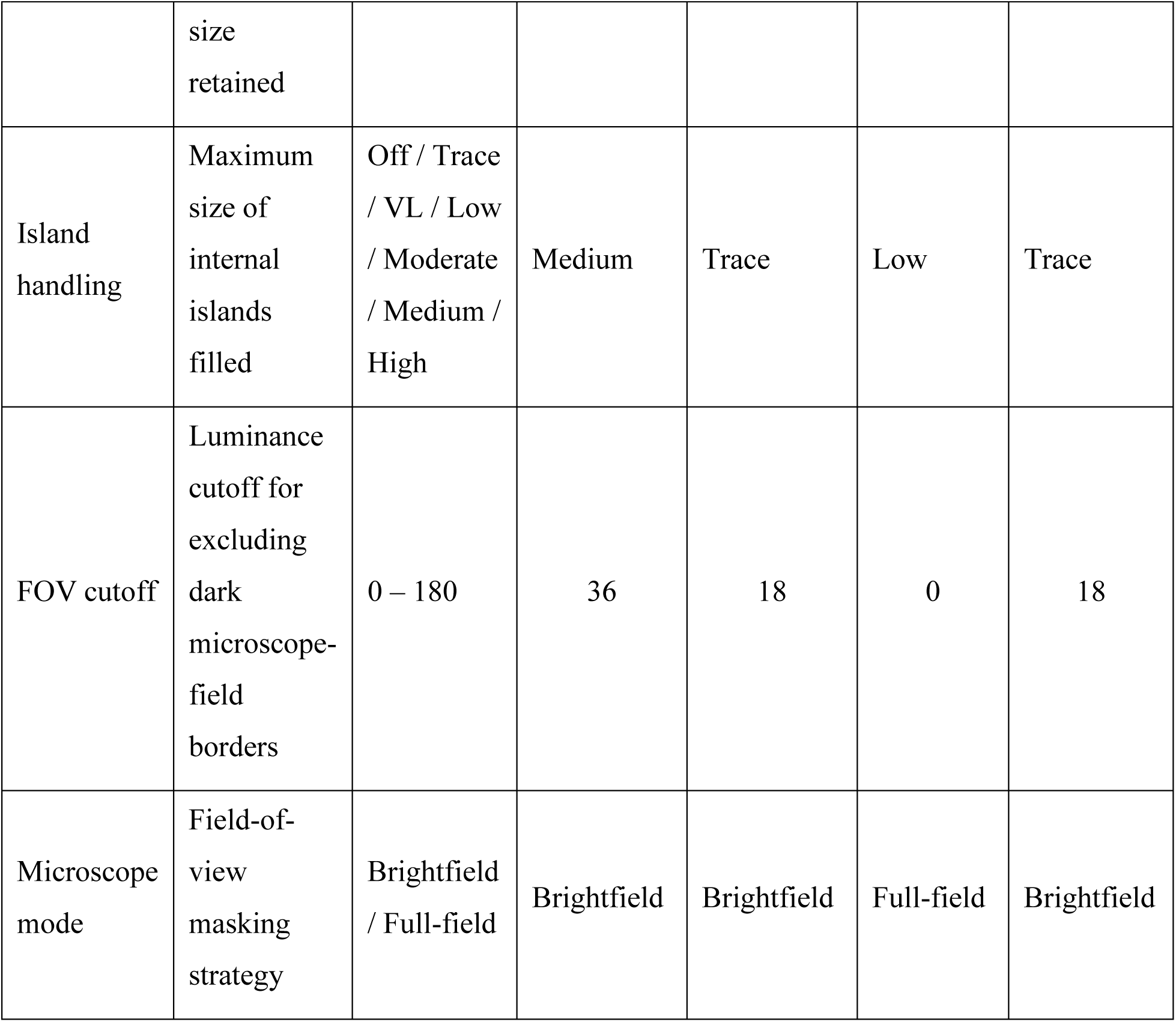
Cytomove segmentation parameters and preset values.

The *Brightfield normal* preset is optimised for standard-contrast brightfield images with clearly delineated wound corridors. The *Brightfield small* preset uses a narrower variance window and aggressive threshold to handle densely packed small-cell monolayers. The *Phase contrast* preset uses full-field masking and a wider variance window suited to phase-contrast images where the cell sheet appears bright-halo. The *Phase contrast 2* preset uses brightfield-style low-variance settings effective for speckled or high-debris phase-contrast datasets.

### Software functionalities

Cytomove integrates adjustable segmentation parameters, overlay-based wound review, quality-control summaries, and frame-level group navigation in a single workspace (Figure 1). It supports two analysis modes. In **single-image mode**, one image is displayed in a review canvas with its segmentation overlay, current measurements, and quality-control labels; the user can adjust parameters and apply manual corrections before exporting. In **group mode**, multiple images (a time course or treatment set) are loaded together, reviewed frame by frame through thumbnail cards, and exported as group-level tables, plots, and overlay archives; each image retains its own segmentation settings stored in memory for the session.

**Figure 1.**
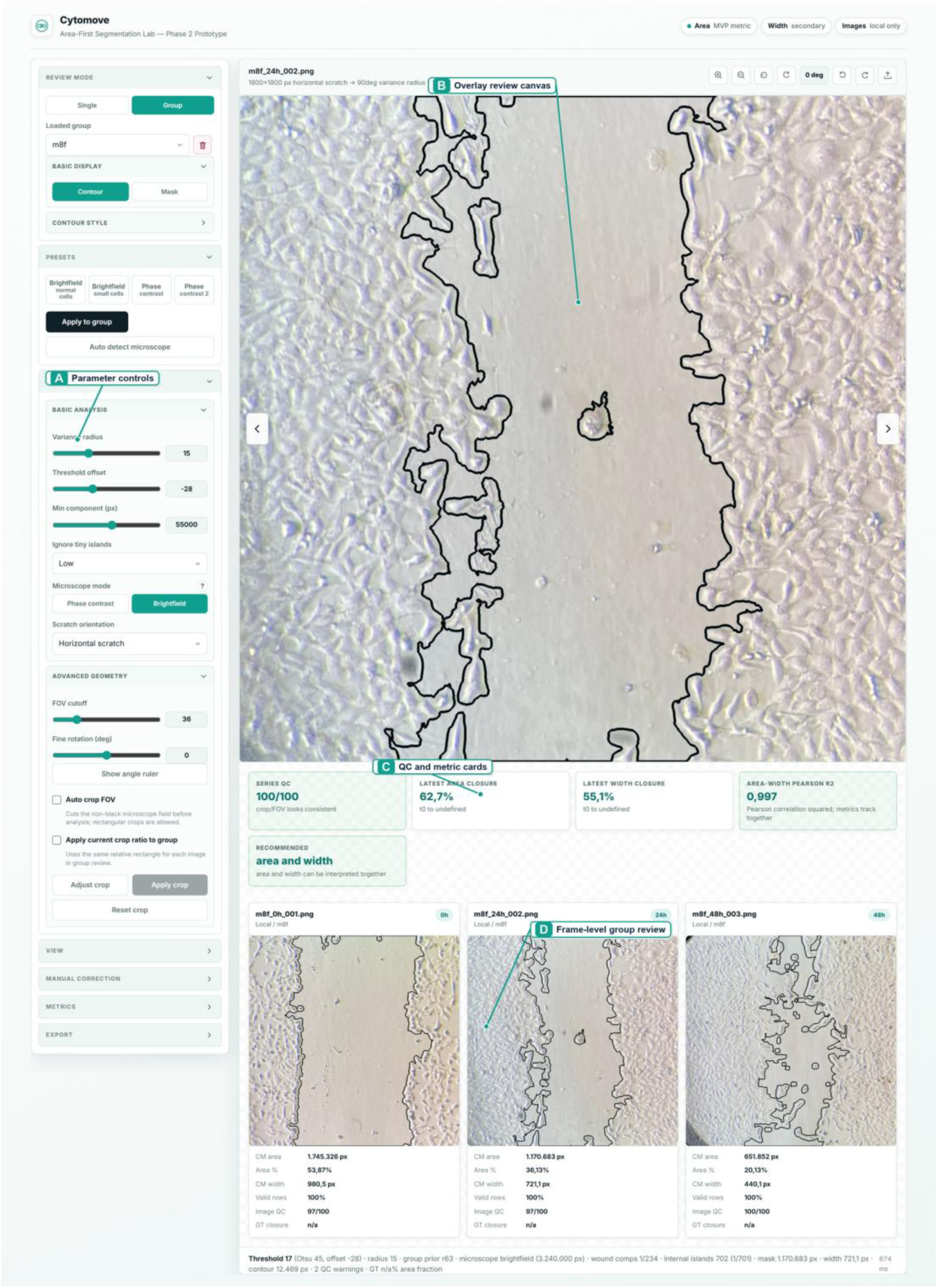
Cytomove application workspace. The interface integrates adjustable segmentation parameters (left panel), overlay-based wound review (centre canvas), quality-control and metric summaries, and frame-level group navigation in a single workflow before export.

The segmentation output is displayed as a contour overlay on the source image rather than as a separate binary mask (Figure 2), letting the user inspect, before exporting any number, whether the wound corridor was captured correctly, whether cells inside the wound were treated appropriately, whether circular field borders were excluded, and whether near-closure regions remain biologically meaningful. When automatic segmentation is insufficient, Cytomove provides four local manual-correction tools that operate on a dragged rectangular region of interest: **Add scan** (local fine threshold, writes back the largest connected gap component); **Fill** (direct region fill); **Erase** (clear mask in region); and **Clean specks** (remove small mask components below a size threshold). Corrections persist across image navigation and are flagged in the export metadata.

**Figure 2.**
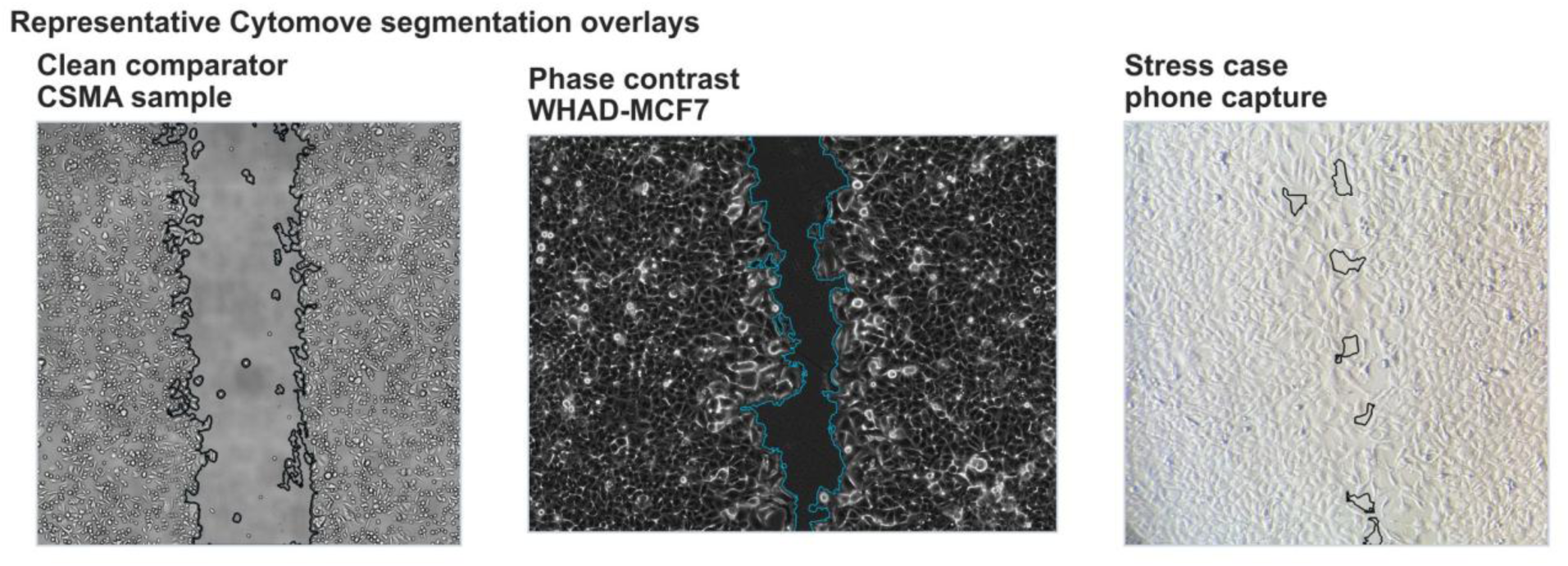
Representative Cytomove segmentation overlays. Contour overlays for (A) a clean brightfield CSMA sample frame, (B) a WHAD-MCF7 phase-contrast frame at mid-closure, and (C) an author-acquired smartphone brightfield frame with circular field borders. The wound contour remains visible so that segmentation quality can be inspected before any measurement is exported.

For every analysed image, Cytomove exports: wound area in pixels; wound area fraction (wound area / field area); mean, median, standard deviation, and coefficient of variation of row-wise wound width; valid-row count and fraction; quality-control labels and warnings; the recommended primary metric (area_and_width, width_preferred, or review_required); crop coordinates and rotation angle; the full set of segmentation parameters; manual-correction status; image dimensions; the algorithm version string; and source metadata where available. Group exports retain the per-frame time point, area, and width values needed to reproduce closure calculations. Exports are provided as CSV and Excel tables (per-image and group-level), PNG overlay images, grouped overlay ZIP archives, and time-course plot images.

### Wound closure percentage

Wound closure percentage at time point t relative to a reference time point t₀ is computed as:

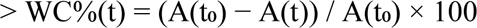

where A(t) is the wound area in pixels at time t. In group review, the earliest or baseline frame is used as the reference for time-course summaries, and the exported per-frame area values allow WC% to be recomputed from the same measurements. For the validation summaries reported here, closure estimates were interpreted descriptively alongside the raw area trajectories and visual overlays rather than treated as independent primary accuracy endpoints.

### Statistical analysis

Agreement between Cytomove and WHST measurements was quantified using five complementary statistics for both wound area and wound width. Let y_i denote the WHST reference value and ŷ_i the Cytomove measurement for paired observation i across n frames in a dataset.

The **mean absolute percentage error** (MAPE) captures average relative discrepancy:

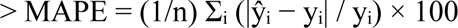

The **median absolute percentage error** is reported alongside MAPE because, in time courses that approach closure, the residual wound becomes very small and a single large relative error can dominate the dataset mean. The **maximum absolute percentage error** identifies the most discrepant frame and guides targeted visual review. The **mean signed percentage error** indicates systematic over- or under-measurement. The **Pearson product-moment correlation coefficient** r captures trend concordance across the time course.

Bland–Altman analysis (Martin Bland and Altman, 1986) was performed on the pooled absolute area scale (in megapixels) rather than as a percentage difference, because percentage differences are numerically unstable for near-closure frames where the denominator approaches zero. The median Cytomove-minus-WHST difference and the 2.5th–97.5th percentile limits of agreement are reported. A non-parametric percentile interval is used rather than ±1.96 SD because the pooled data include time-course frames that are not independent observations. All statistics were computed in Python using NumPy and SciPy.

### Validation datasets and comparator

Cytomove was evaluated against WHST across five image sets comprising 31 paired measurements (Table 2). WHST was selected as the primary quantitative benchmark because its ImageJ/Fiji output vocabulary (wound area, wound area fraction, average width, width standard deviation) overlaps directly with Cytomove’s exported endpoints. For each dataset, signed and absolute percentage differences between Cytomove and WHST were computed per frame and summarised at the dataset level.

**Table 2.**
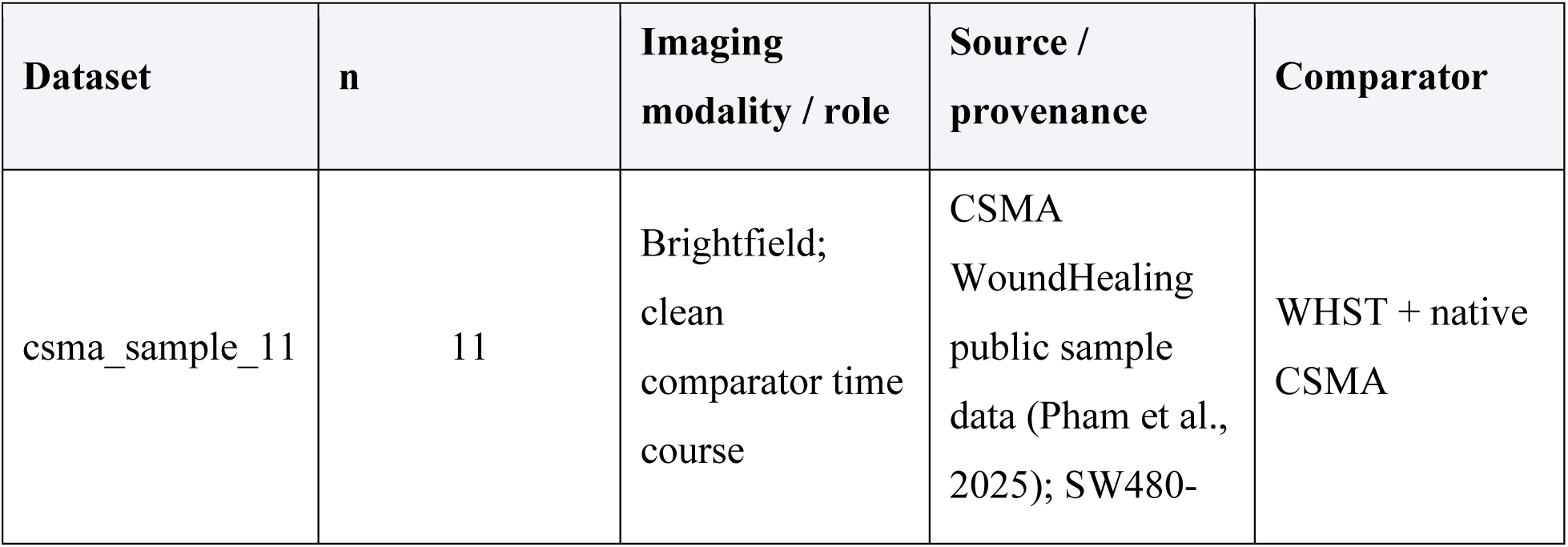
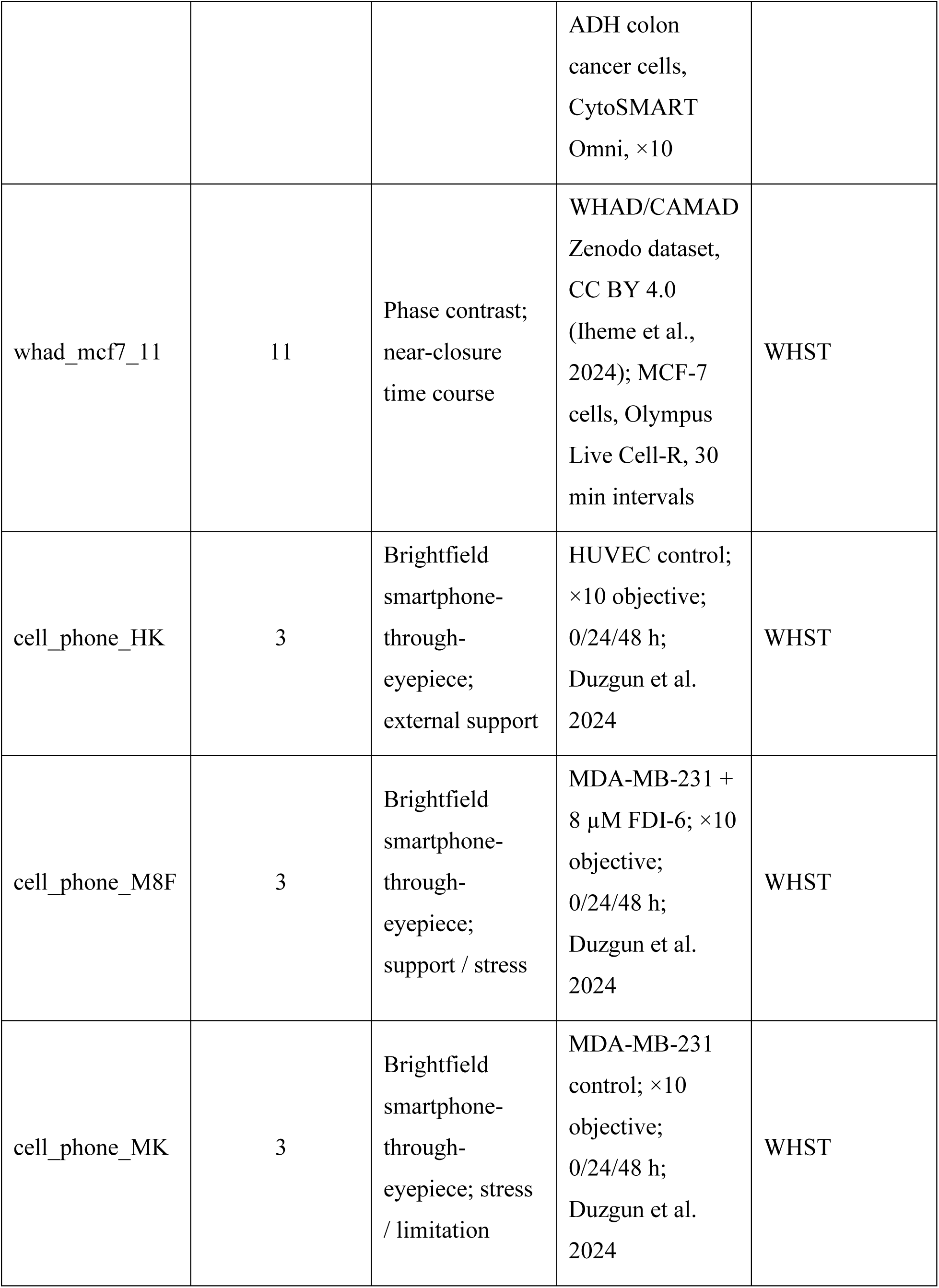
Preliminary validation datasets.

| **Dataset** | **n** | **Imaging modality / role** | **Source / provenance** | **Comparator** |
| --- | --- | --- | --- | --- |
| csma_sample_11 | 11 | Brightfield; clean comparator time course | CSMA WoundHealing public sample data (Pham et al., 2025); SW480- | WHST + native CSMA |
|  |  |  | ADH colon cancer cells, CytoSMART Omni, ×10 |  |
| whad_mcf7_11 | 11 | Phase contrast; near-closure time course | WHAD/CAMAD Zenodo dataset, CC BY 4.0 (Iheme et al., 2024); MCF-7 cells, Olympus Live Cell-R, 30 min intervals | WHST |
| cell_phone_HK | 3 | Brightfield smartphone-through-eyepiece; external support | HUVEC control; ×10 objective; 0/24/48 h; Duzgun et al. 2024 | WHST |
| cell_phone_M8F | 3 | Brightfield smartphone-through-eyepiece; support / stress | MDA-MB-231 + 8 µM FDI-6; ×10 objective; 0/24/48 h; Duzgun et al. 2024 | WHST |
| cell_phone_MK | 3 | Brightfield smartphone-through-eyepiece; stress / limitation | MDA-MB-231 control; ×10 objective; 0/24/48 h; Duzgun et al. 2024 | WHST |

**csma_sample_11.** An 11-frame brightfield time-course subset drawn from the CSMA WoundHealing public sample dataset (Thanh Pham et al., 2025). The original sequence comprises SW480-ADH human colon cancer cells imaged every hour for 48 hours at ×10 magnification on an Omni Cell Imager (CytoSMART Technologies, The Netherlands); images are JPEG files cropped to approximately 1537 × 1537 pixels. Eleven representative frames were selected to span the full time course at approximately equal intervals. Native CSMA, WHST, and Cytomove outputs were available for the matched frames, enabling a three-method area comparison.

**whad_mcf7_11.** An 11-frame phase-contrast time-course sequence comprising MCF-7 human breast cancer cells from the publicly deposited Wound Healing Assay Dataset (WHAD) and Cell Adhesion and Motility Assay Dataset (CAMAD) (Iheme et al., 2024), released under Creative Commons Attribution 4.0 International. The sequence was acquired on an Olympus Live Cell-R Station at 30-minute intervals; cells approach near-complete closure by the final frames, making this set the primary test of Cytomove behaviour in challenging late time-course conditions.

**Author-acquired smartphone-through-eyepiece sets (HK, M8F, MK).** Three brightfield time-course sets (three frames each: 0, 24, and 48 hours) acquired by the author using a smartphone camera held to the eyepiece of an inverted microscope at ×10 magnification. The HK set used Human Umbilical Vein Endothelial Cells (HUVEC; ATCC), whereas the M8F and MK sets used MDA-MB-231 human metastatic breast cancer cells. Cells were grown to confluence in DMEM supplemented with 10% FBS and 1% penicillin–streptomycin at 37°C in 5% CO₂, then scratched with a 1 mL pipette tip in a 6-well plate and imaged at 0, 24, and 48 hours. The M8F set was treated with 8 µM FDI-6 (Sigma-Aldrich, cat. no. SML1392-5MG) for the duration of the time course; HK and MK are untreated controls from the same experimental series. This experimental series was originally reported in (Duzgun et al., 2025). The images were acquired without a motorised stage, which produced variable circular field borders, uneven illumination, and focus variation similar to the acquisition conditions encountered in many laboratories. These sets are included in the validation as external support and stress material, not as primary accuracy evidence.

## Results

The validation was designed to ask a practical question: whether a browser-local, reviewable implementation can reproduce the wound-area behaviour of an established ImageJ/Fiji comparator while making difficult frames visible to the user. The five sets combine a clean public brightfield comparator sequence, a public phase-contrast time course that approaches closure, and three author-acquired smartphone sets used as external support or stress material. The numerical summary is in Table 3, and each main result is paired with visual overlays or audit panels in Figures 2–6.

**Figure 3.**
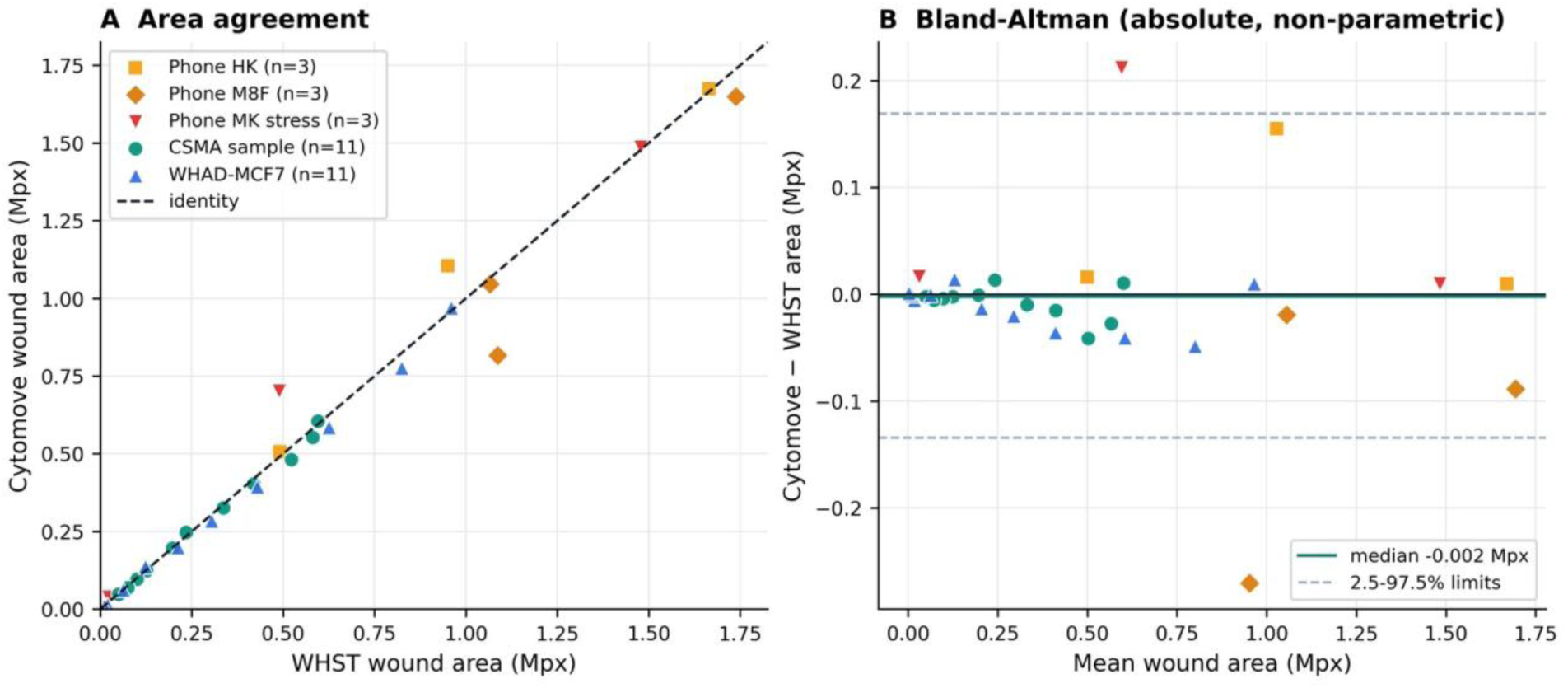
WHST versus Cytomove wound-area comparison across all validation sets. (A) Paired WHST and Cytomove wound-area values against the identity line, with a distinct marker per dataset. (B) Bland–Altman view on the absolute area scale; the solid line marks the median difference (−0.002 Mpx) and the dashed lines the 2.5–97.5th percentile limits of agreement (−0.134 to +0.169 Mpx).

**Figure 4.**
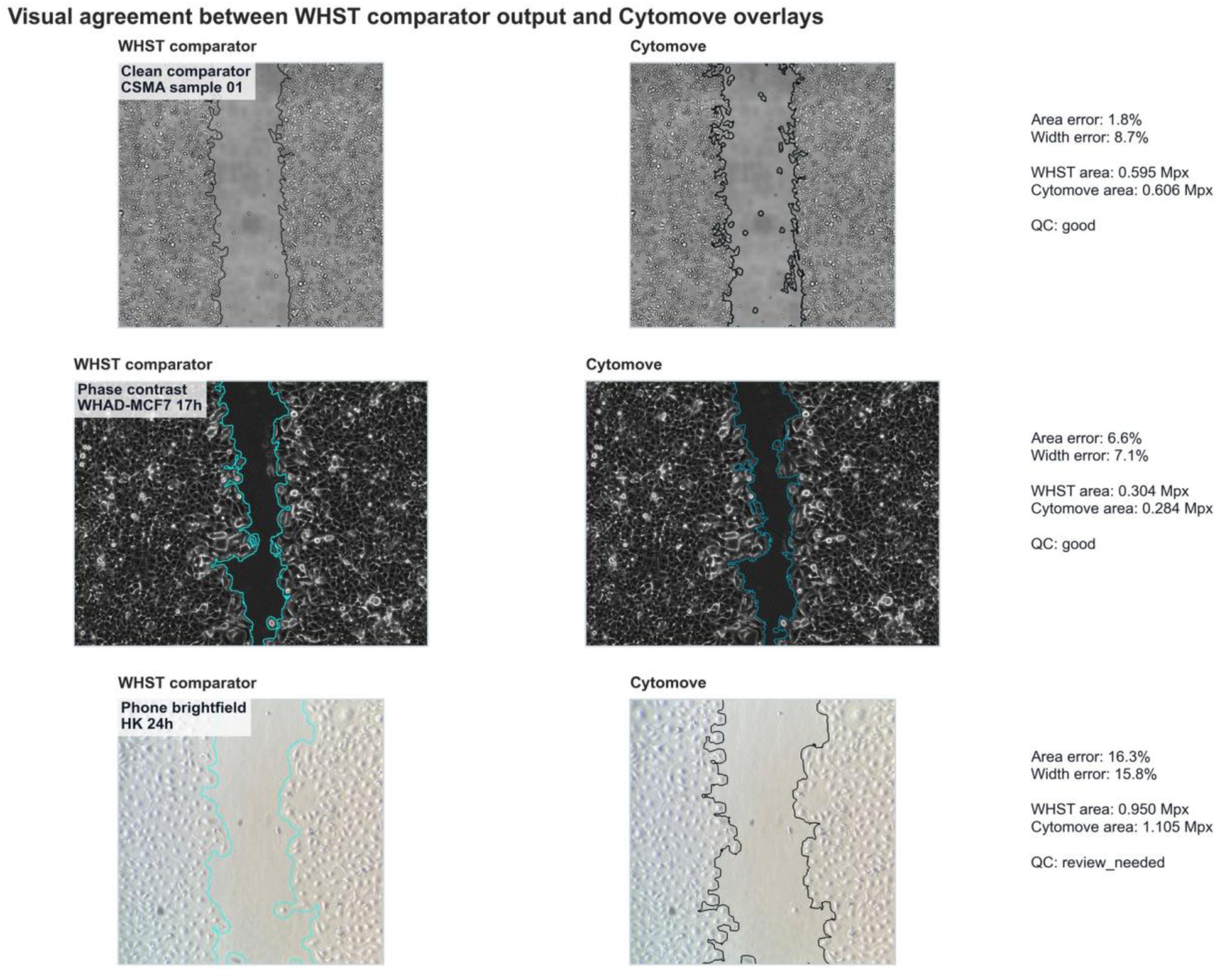
Visual comparison of WHST output and Cytomove overlays. Representative frame pairs connecting numerical comparison with visual evidence across the clean brightfield, phase-contrast, and smartphone acquisition contexts.

**Figure 5.**
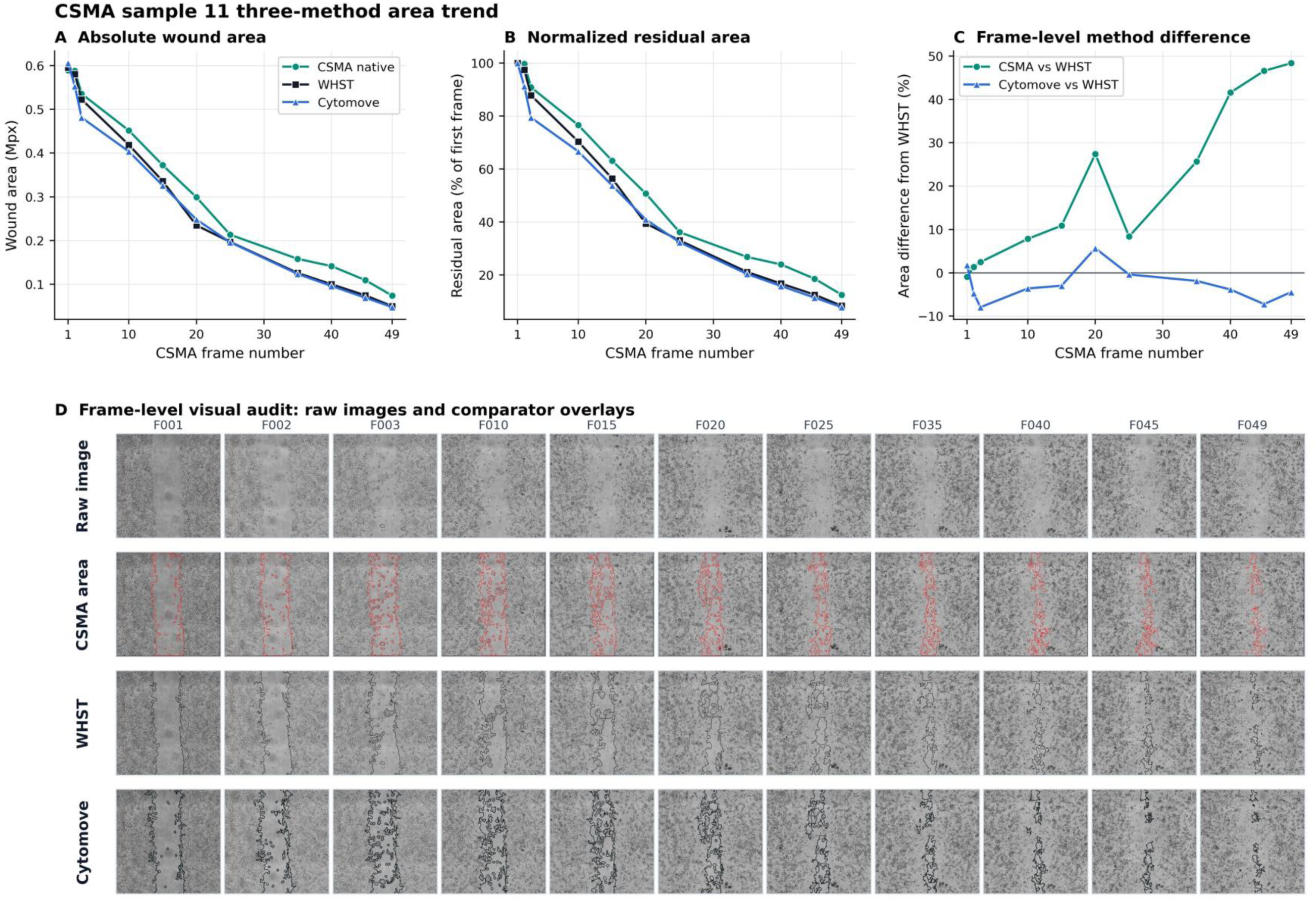
CSMA sample 11 three-method area comparison and frame-level visual audit. (A–C) Native CSMA, WHST, and Cytomove wound area across the 11-frame sequence; (D) frame-level audit showing raw image, CSMA area output, WHST measurement, and Cytomove overlay for each frame.

**Figure 6.**
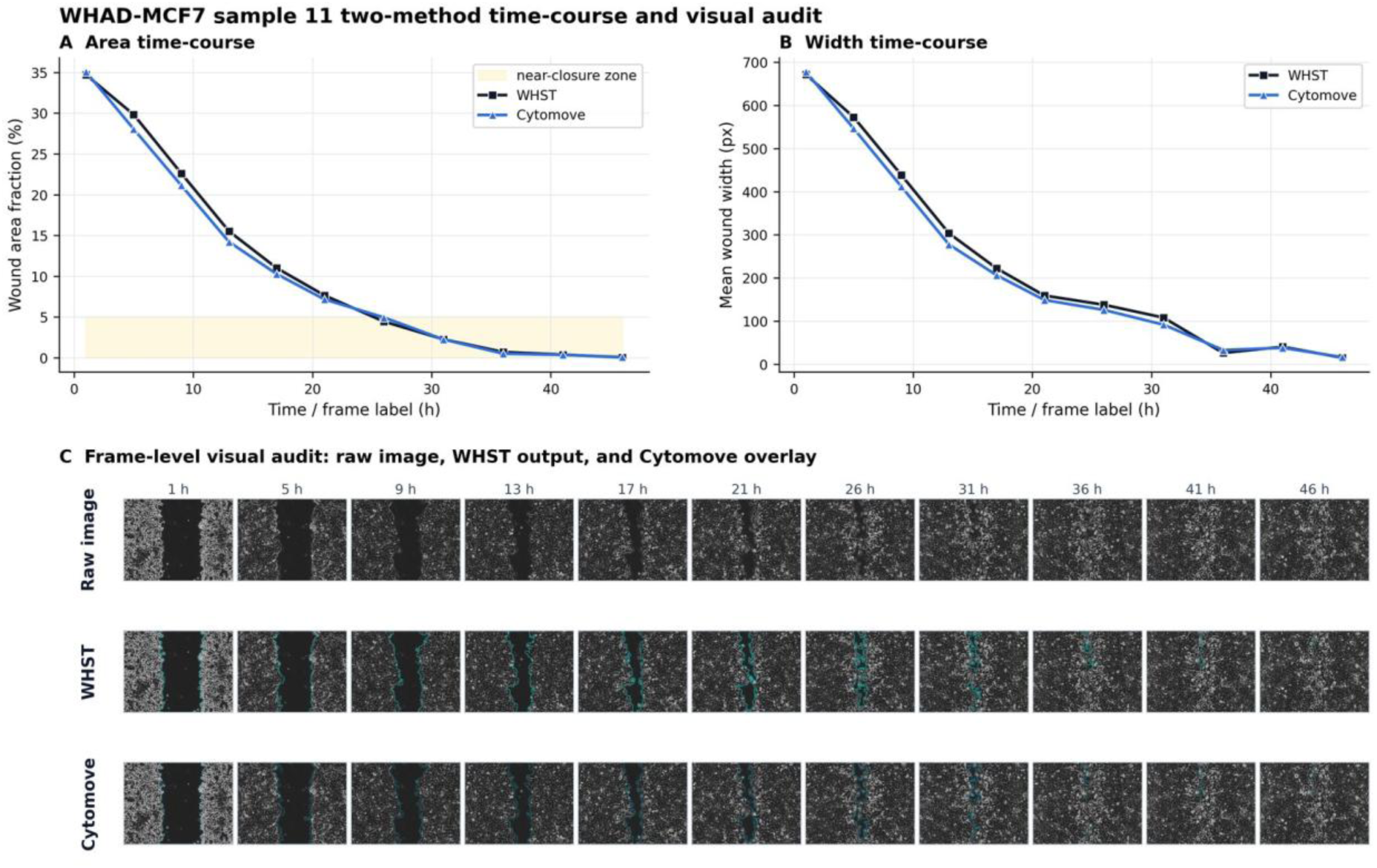
WHAD-MCF7 phase-contrast time-course and frame-level audit. (A) Wound area and (B) wound width trajectories across the 11-frame sequence, with (C) frame-level overlays; late near-closure frames are flagged as review-sensitive.

**Table 3.** Preliminary Cytomove-versus-WHST agreement summary. Area and width are reported as mean absolute percentage error (MAPE), median absolute percentage error, maximum absolute percentage error, mean signed percentage error, and Pearson trend correlation (r).

| Dataset | n | Area<br>MAPE % | Area<br>median % | Area<br>max % | Area<br>signed % | Area<br>r | Width<br>MAPE % | Width<br>r | Interpretation |
| --- | --- | --- | --- | --- | --- | --- | --- | --- | --- |
| csma_sample_11 | 11 | 4.1 | 3.9 | 7.9 | -2.7 | 0.9975 | 9.7 | 0.9969 | Primary clean comparator; visually concordant overlays |
| whad_mcf7_11 | 11 | 15.0 | 6.6 | 82.5 | +2.2 | 0.9984 | 9.0 | 0.9986 | Phase contrast; MAPE driven by late near-closure frames |
| cell_phone_HK | 3 | 6.7 | 3.2 | 16.3 | +6.7 | 0.9903 | 8.5 | 0.9918 | External support set (HUVEC) |
| cell_phone_M8F | 3 | 10.6 | 5.1 | 24.9 | -10.6 | 0.9557 | 8.7 | 0.9076 | Support / stress; late-frame underestimation in FDI-6- |
|  |  |  |  |  |  |  |  |  | treated MDA-MB-231 |
| cell_phone_MK | 3 | 38.8 | 43.4 | 72.3 | +38.8 | 0.9880 | 31.0 | 0.9861 | Stress / limitation case; fragmented near-closed wound; not accuracy evidence |

**Table 4.** Software and code metadata.

| Nr. | Code metadata description | Value |
| --- | --- | --- |
| C1 | Current code version | v1.0.0 |
| C2 | Permanent link to code/repository | https://github.com/zduzgun/CytoMove (public project repository) |
| C3 | Permanent link to Reproducible Capsule | Not applicable |
| C4 | Legal Code License | Source-available non-commercial licence following the PolyForm Noncommercial License 1.0.0 model; commercial use is not permitted under the public licence and requires a separate written commercial licence |
| C5 | Code versioning system used | git |
| C6 | Software languages, tools, and services | JavaScript (ES2020+), HTML5 Canvas API, typed arrays (Uint8Array, Float32Array, Int32Array); Electron (desktop package) |
| C7 | Compilation requirements and dependencies | Runs in any modern Chromium- or Firefox-based browser with no installation or dependency; desktop build requires Node.js $\geq 18$ and Electron |
| C8 | Link to developer documentation | https://github.com/zduzgun/CytoMove (README and /docs) |
| C9 | Support email |  |

### Clean brightfield comparator: csma_sample_11

The csma_sample_11 subset provided the clearest support for Cytomove accuracy. Wound-area measurements agreed closely with WHST (area MAPE 4.1%, median 3.9%, maximum 7.9%, mean signed error −2.7%, Pearson r = 0.9975); width agreement was acceptable (MAPE 9.7%, r = 0.9969). Wound closure percentage tracked the expected monotonic increase from 0% at frame 1 to approximately 85–90% at frame 11, consistent with the known behaviour of SW480-ADH cells in the CSMA sample series.

A three-method comparison was possible because native CSMA, WHST, and Cytomove area outputs were available for the matched frames (Figure 5). All three followed the same decreasing wound-area trajectory across the sequence (CSMA–WHST r = 0.9970; CSMA–Cytomove r = 0.9951). The main difference was not the direction of the trajectory but the residual area retained by native CSMA in later frames, a pattern consistent with CSMA’s two-stage wound-edge and cell-detection approach, which can attribute some cell-free space differently from a pure low-variance mask. This difference is presented as method-context evidence rather than a failure case: different tools can preserve the same biological trend while making different boundary choices, and Cytomove’s overlay makes those choices inspectable.

The visual audit panel in Figure 5D shows raw images alongside CSMA area output, WHST measurement overlay, and Cytomove contour for each frame. In early frames (large open wound), all three methods produce visually concordant boundaries; differences become apparent only at late frames when the wound narrows below approximately 100–150 pixels in width.

### Phase-contrast time course: whad_mcf7_11

The whad_mcf7_11 set tested Cytomove on a wound that progresses to near-complete closure (Figure 6). Cytomove followed the WHST area and width trends closely (r = 0.9984 and r = 0.9986) with a modest median absolute area error of 6.6% and a small positive mean signed error (+2.2%). The dataset-level area MAPE of 15.0% was concentrated in the latest near-closed frames: in the final frame the residual wound measured approximately 1,400 pixels by WHST versus approximately 2,550 pixels by Cytomove, an absolute difference of roughly 1,150 pixels that registers as an ∼82% relative error due to the very small denominator.

Cytomove flags such frames with quality-control labels (review_needed, area_ok_width_review) and exposes the overlay, valid-row information, and width variability coefficient in the export so that the user can decide whether the measurement is acceptable before reporting it. The median error of 6.6% gives a more representative summary of the 11-frame sequence than the mean, and the maximum identifies the single frame requiring closest attention. Wound closure percentage computed by Cytomove for this sequence agreed with the WHST-derived trajectory in the biologically informative early and mid time-course phases; divergence appeared only in the final 1–2 frames where both tools measure a very small residual wound.

### Smartphone-through-eyepiece stress sets: HK, M8F, MK

Three smartphone-through-eyepiece brightfield sets reproduce conditions common in real laboratories but absent from curated public datasets: circular fields, uneven illumination, focus variation, and fragmented late wounds. Results are summarised in Table 3 and Figure 4.

The HK (HUVEC control) set behaved as a supportive external example (area MAPE 6.7%, r = 0.9903). The M8F set (MDA-MB-231 treated with 8 µM FDI-6) showed a late-frame underestimation pattern (area MAPE 10.6%, mean signed error −10.6%), consistent with the fragmented wound appearance at 48 h when FDI-6 partially suppresses MDA-MB-231 cell migration (Duzgun et al., 2025). The MK (MDA-MB-231 control) set behaved as a genuine limitation case: Cytomove over-measured fragmented near-closed regions (area MAPE 38.8%, width MAPE 31.0%), with every difficult frame flagged for review by the quality-control system. Wound closure percentage estimates for HK and MK reached approximately 70–80% at 48 h; M8F reached approximately 55–65%, consistent with the treatment effect reported in the original study.

These sets are reported as quality-control and stress examples. Their value lies in showing that Cytomove can identify frames requiring user inspection, preserve the overlay evidence for that decision, and record the correction or acceptance in the export metadata, not in claiming solved accuracy under all acquisition conditions.

### Pooled agreement and Bland–Altman analysis

A pooled Bland–Altman comparison on the absolute area scale (Figure 3B) showed a median Cytomove-minus-WHST difference of essentially zero (−0.002 Mpx), with 2.5–97.5th percentile limits of agreement of −0.134 to +0.169 Mpx. The paired area plot (Figure 3A) shows that most measurements fall close to the identity line across the clean and phase-contrast sequences; the phone-capture limitation set (MK) contributes the more visibly dispersed points. Area was the more stable and comparable metric across all sets; width tracked area trends but was more sensitive in fragmented and near-closed frames (Figure 4; Supplementary Figure S1).

## Discussion

### Main finding

The main finding is that Cytomove can reproduce WHST-like wound-area behaviour in image sets that are visually suitable for variance-and-threshold segmentation, while preserving enough visual and metadata context to identify frames that should not be interpreted automatically. The clean brightfield CSMA sequence provides the strongest agreement evidence. The WHAD-MCF7 phase-contrast sequence supports the same direction but also demonstrates why near-closure frames need careful reporting: small residual wounds turn modest absolute differences into large percentage errors. The smartphone sets broaden the evaluation by showing how Cytomove’s quality-control workflow responds to real-world acquisition challenges.

### Workflow and reproducibility contribution

Cytomove is a workflow and reproducibility contribution rather than a new biological metric. The established scratch-assay literature already defines the major measurement concepts (wound area, normalised area, width, closure percentage, time-course plots, output images, and spreadsheet export) across TScratch, WHST, PyScratch, and CSMA. Cytomove brings these into a browser-local, reviewable workflow that reduces installation friction while preserving inspectability and recording the provenance of each measurement.

Specifically, every Cytomove export includes the image identifier, crop coordinates, scratch orientation, fine rotation angle, variance radius, threshold offset, minimum component size, island-handling level, field-of-view cutoff, microscope mode, manual-correction status, algorithm version string, and quality-control labels. To reproduce a measurement, a researcher needs all of these, and Cytomove records them automatically in a machine-readable CSV or Excel file. This directly addresses the common loss of analysis settings in macro- and plugin-based workflows.

### Positioning relative to existing tools

Cytomove shares the core design lesson of TScratch, that automated segmentation should remain visually inspectable and manually editable, while implementing it in a browser-local architecture. It shares WHST’s measurement vocabulary (area, area fraction, average width, width standard deviation) and extends it with a richer export schema. It shares PyScratch’s emphasis on usability for non-programmer researchers, but instead of a Python GUI application it requires only a modern browser and no installation step. It addresses the same wound-internal cell concern that motivated CSMA, though through a user-inspectable overlay and manual-correction tools rather than a two-stage automated cell-detection pipeline.

The absence of a server-side backend is a deliberate design decision. Browser-local analysis eliminates the privacy concern associated with uploading unpublished microscopy to a remote service and removes the operational cost and availability risk of a server. The JavaScript implementation imposes a computational limit relative to native Python or Java backends, but for the image sizes and parameter ranges used in typical scratch assays, the typed-array pipeline runs well within the five-second-per-image target on consumer hardware.

### Near-closure frames and metric choice

The treatment of near-closure frames deserves particular caution. In a wound-healing time course, the final biological state may be the most interesting, but it is also where relative error becomes least stable. A difference of roughly a thousand pixels can be small in the context of early open wounds yet large as a fraction of a nearly closed residual gap. This is why reporting MAPE, median error, maximum error, and Pearson r together is essential: correlation captures trend agreement, MAPE captures average relative discrepancy, median error reduces the influence of the near-closure tail, and the maximum identifies the frame a user should inspect. Area is treated as the primary endpoint because it was more stable across the current validation sets; width serves as a secondary quality-control endpoint that is more sensitive to fragmentation and partial closure.

### Limitations and planned work

The present validation should be interpreted as preliminary. The strongest evidence comes from the csma_sample_11 and whad_mcf7_11 sequences; the smartphone sets are included to expose review and quality-control behaviour rather than to claim solved accuracy. The study does not yet include multi-rater manual consensus masks, Dice coefficient or intersection-over-union evaluation (Maier-Hein et al., 2024; Taha and Hanbury, 2015), spatial calibration for physical-unit closure rates, runtime benchmarking across browsers and operating systems, wound-internal cell detection (Thanh Pham et al., 2025), or a systematic cross-browser reproducibility test.

Recent bioimage-analysis literature shows how rapidly segmentation methods are advancing. Generalist and foundation-model approaches such as Cellpose3, Segment Anything for Microscopy, and MedSAM have demonstrated impressive segmentation generalisation (Archit et al., 2025; Ma et al., 2024; Stringer and Pachitariu, 2025). An explainable local pipeline with visible overlays can remain useful alongside these methods because the workflow problem is not only segmentation accuracy but also review, provenance, export, and reproducible reporting. Any future learned-segmentation backend in Cytomove should be optional, reviewable, and evaluated against the same metadata and visual-audit standards used here.

Planned work includes: multi-rater consensus mask validation with spatial-overlap metrics; spatial and temporal calibration for physical-unit area and migration rate; wound-internal cell handling; runtime and cross-browser reproducibility testing; and export packages aligned with FAIR data principles (Wilkinson et al., 2016).

### Conclusion

Cytomove provides a browser-local and reviewable workflow for scratch wound healing assay quantification. It implements an explainable variance-and-threshold pipeline in dependency-free JavaScript, supports single-image and grouped time-course analysis, and exports a complete audit trail linking every measurement to the segmentation image, parameters, and quality-control state that produced it. The current evidence supports Cytomove as an accessible, publication-oriented analysis tool that can produce WHST-comparable wound-area outputs in suitable brightfield and phase-contrast images, while preserving the overlay and quality-control context needed to interpret near-closure and real-world acquisition challenges.

## Ethics statement

This study did not involve human participants, identifiable human data, or animal experiments, and therefore required no ethical approval. The MCF-7 phase-contrast frames were obtained from the publicly available WHAD/CAMAD dataset released under the Creative Commons Attribution 4.0 International licence (Iheme et al., 2024). The CSMA brightfield sample frames and output context were obtained from the public CSMA WoundHealing repository associated with (Thanh Pham et al., 2025). The smartphone-through-eyepiece brightfield images were acquired from established cell-line cultures as part of the study described in (Duzgun et al., 2025).

## CRediT author statement

**Zekeriya Duzgun:** Conceptualization, Methodology, Software, Validation, Formal analysis, Data curation, Visualization, Writing – original draft, Writing – review & editing.

## AI assistance statement

The author used AI-assisted tools during manuscript preparation for language editing, formatting support, and workflow organization. The author reviewed and verified all scientific content, analyses, interpretations, references, and final text, and takes full responsibility for the manuscript.

## Data and software availability

A live web version is available at https://cytomove.com, and the public project repository is https://github.com/zduzgun/CytoMove. The core analysis source code is intended to be publicly available under a source-available non-commercial licence, following the PolyForm Noncommercial License 1.0.0 model; commercial use is not permitted under the public licence and requires a separate written commercial licence from the author or rights holder. The preliminary validation summary (paired WHST-Cytomove measurements, dataset-level diagnostic statistics, the discrepancy-resolution table, selected input provenance, derived measurement outputs, and representative overlay images) is available from Zenodo at https://doi.org/10.5281/zenodo.20486820. External image provenance: WHAD-MCF7 frames from the WHAD/CAMAD Zenodo dataset under CC BY 4.0 (Iheme et al., 2024); CSMA sample images and output context from the public CSMA WoundHealing repository (Thanh Pham et al., 2025); smartphone-through-eyepiece images acquired by the author as part of the experimental series reported in (Duzgun et al., 2025).

---

**Supplementary Figure S1.**
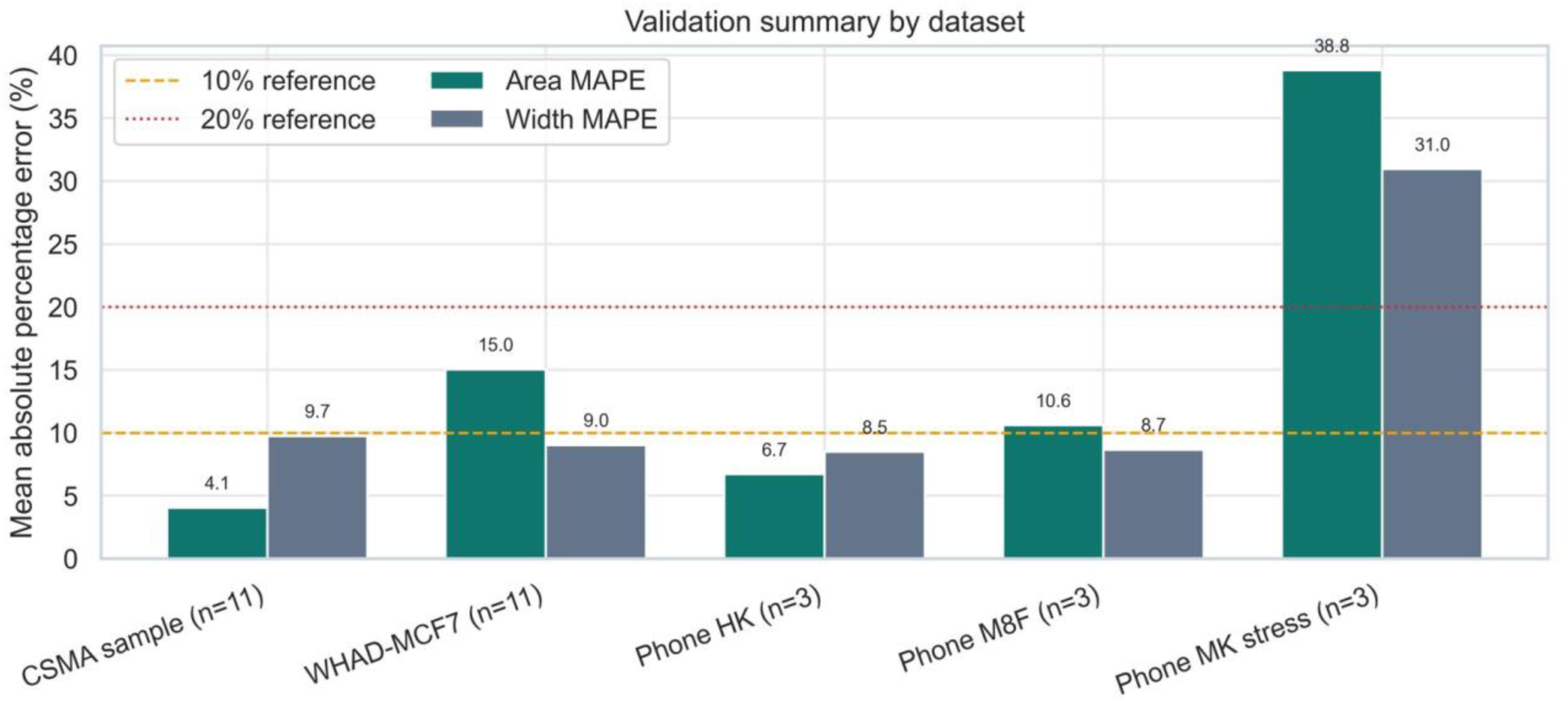
Dataset-level validation summary. Diagnostic panel of area and width agreement statistics across all five validation sets.

