## Supplementary figures and images for "Cytomove: a browser-local and reviewable workflow for scratch wound healing assay quantification"

### Supplementary Figure S1

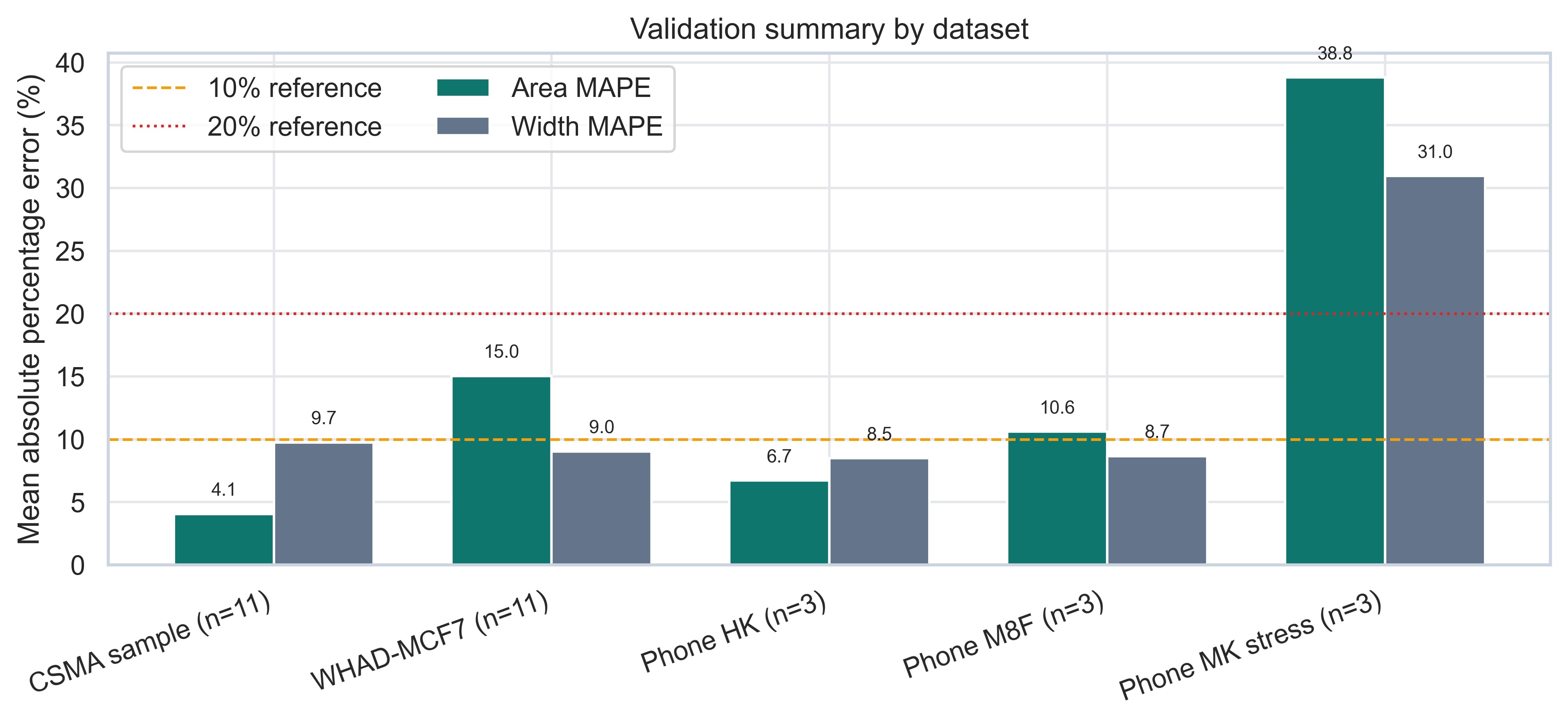
